# Hippocampus serves as a repository for spoken and heard word meanings during conversations

**DOI:** 10.1101/2025.09.20.677504

**Authors:** Ana G. Chavez, Xinyuan Yan, Melissa Franch, Elizabeth A. Mickiewicz, Will Baltazar, James L. Belanger, Davin Devara, Maya Etta, Thomas Hamre, Taha Ismail, Brad Joiner, Yoon Kim, Abhinav Kona, Kasra Mansourian, Amy Nangia, Molly Pluenneke, Sarah Soubra, Thara Venkateswaran, Karthik Venkudusamy, Assia Chericoni, Katherine E. Kabotyanski, Kalman A. Katlowitz, Raissa K. Mathura, Tarana Nigam, Danika L. Paulo, Hanlin Zhu, Eleonora Bartoli, Nicole R. Provenza, Andrew Watrous, Krešimir Josić, Sameer A. Sheth, Benjamin Y. Hayden

**Affiliations:** Neurosurgery, Baylor College of Medicine; Neuroengineering Initiative, Rice University; Departments of Mathematics, Biology and Biochemistry, University of Houston

## Abstract

We utilize internal representations of meaning for two purposes: to understand the words we hear and to generate our own speech. This dual requirement necessitates abstract, modality-agnostic representations. Building on work identifying it as a hub for relational mapping, we hypothesized that the hippocampus supports abstract, cross-person representations, and uses shared semantic geometries to do so. We tested this hypothesis by examining hippocampal activity in a remarkable single-neuron dataset derived from conversational speech. Neurons robustly encoded meanings of both spoken and heard words, and used common geometric embeddings for both, leading to abstract meaning performance. Speaker identity was aligned with meaning via partial subspace alignment, which affords speaker-meaning binding by partitioning meaning by speaker while maintaining cross-speaker generalization. Degrees of subspace rotation varied on a single word level and depended systematically on semantic category. Together, these findings indicate how geometric principles allow for abstract cross-personal meanings while preserving binding to speaker identity.

## INTRODUCTION

Communication relies on speaking and listening, which link form and meaning in different ways. These two processes have distinct sensory and motor expressions and are associated with different goals and mental operations. On the other hand, they make use of a shared abstract representation of meaning. That shared representation of meaning must be accessible to both speaking and listening, so that we can fluidly move back and forth between speaking and listening on common topics. If one person mentions the “*moon*”, they are probably thinking about the same moon as their partner. Identifying how meaning is represented in a manner that affords both processes is an important goal in neuroscience (Menenti et al., 2011).

Classic approaches to the functional neuroanatomy of language emphasize the specialization of regions for language production and reception (such as Broca’s and Wernicke’s Areas) but cannot readily account for cross-personal generalization of meaning. More modern approaches suggest that there are anatomical overlaps between the systems used for speaking and listening, but do not explain how this alignment occurs computationally (Cogan et al., 2014; Silbert et al., 2014). One potential solution to this problem comes in the form of specialized mirror neurons whose responses encode meanings regardless of speaker (Arbib, 2005; Corballis, 2010; Rizzolati and Arbib, 1998; Rizzolatti and Craighero, 2004). However, the functional interpretation of such responses remains debated, as sensorimotor activations during speech perception may reflect downstream or modulatory processes rather than the substrate of semantic understanding itself (Heyes and Catmur, 2022; Hickok, 2010 and 2014, Toni et al., 2008). This perspective suggests that semantic meaning is most appropriately studied at the level of distributed conceptual representations, which can support the flexible recombination and graded distinctions in meaning observed in natural language (Ebitz and Hayden, 2016; Quiroga et al., 2020; Rigotti et al., 2016).

Recent advances in the geometry of representation raise an alternative possibility (Barak et al., 2013; Bernardi et al., 2020; Johnston et al., 2024; Perich et al., 2025). Word meanings can be represented in the geometric organization of encoding manifolds (Franch et al., 2025; Yan et al., 2025). These codes can be aligned to serve different purposes - like speaking and listening - without losing their overall structure. We are particularly interested in the hippocampus due to its demonstrated role in conceptual representations, as well as in generalization and relational reasoning (Bakermans et al, 2025; Bakker et al., 2008; Bernardi et al. 2020; Courellis et al., 2024; Leutgeb et al., 2005). Hippocampus has a well-established role in language, especially in semantic representations (Binder et al., 2009; Dijksterhuis et al., 2024; Duff and Brown-Schmidt, 2012; Franch et al., 2025; Katlowitz et al., 2025; Piai et al., 2016; Van de Ven et al., 2020), and its clear role in identity assignment (Danjo et al., 2018; Montagrin et al., 2018; Rey et al., 2020; Tavares et al., 2015). These representations appear to emerge from the hippocampal extraction of latent relational structures, enabling the formation of compositional schemas that transcend specific use cases (Bakermans et al., 2025; Bernard et al., 2026). By instantiating high-dimensional manifolds that disentangle disparate features of experience, the hippocampus may provide the geometric flexibility necessary to align knowledge across distinct modalities and symbolic formats without sacrificing representational specificity (Tang et al., 2025; Whittington et al., 2022). These findings endorse the hypothesis that the hippocampus serves as a general repository of meaning that can be accessed by both comprehension and production systems.

We analyzed speech data with a unique dataset collected during natural bidirectional communication - unconstrained conversations. This design makes cross-speaker semantic alignment inherently harder to detect, but provides a more ethologically valid test of whether shared representations of meaning arise during real-world language use. We studied patients in the epilepsy monitoring unit (EMU) conversing with members of their family or our research team. We manually transcribed all spoken words and computed their semantic embeddings using BERT. We aligned each word to firing rates of neurons in the hippocampus and used encoding models to estimate semantic tuning (Franch et al., 2025). Encoding models are particularly powerful because they allow us to limit the features of interest to lexicon, and control for factors like intonation, pitch, and so forth. We found robust encoding of lexical semantics for both production and reception, with strongest encoding around the time of utterance for production, and immediately after it for reception. Critically, we find shared geometric representation of semantics for speaking and for listening and, moreover, evidence for cross-conditional generalization for semantic categories (Bernardi et al., 2020; Kriegeskorte and Wei, 2021). Moreover, that speaking and listening occupy partially overlapping subspaces, affording speaker-specific differentiation without sacrificing cross-speaker generalization of meaning (Johnston et al., 2024). Finally, the degree of overlap varied consistently with semantic category: categories with the greatest generalization included less personalized words, such as function words and pronouns; categories with greatest differentiation were more specifically personalized, such as body parts and proper nouns. Together these findings highlight the power of geometrical embeddings for reconciling the dual demands of speaking and listening while maintaining an abstract representation of meaning.

## RESULTS

Ten native English-speaking patients undergoing neural monitoring for epilepsy participated in unstructured conversations in the epilepsy monitoring unit (EMU) at Baylor St. Luke’s Hospital. All patients had multiple Behnke-Fried electrodes placed in the hippocampus for surgical monitoring (**Figure 1A**). Word counts across all conversations spanned from 720 to 8260 total words (mean: 5267 words). Conversations involved either two (n=3/10) or three-plus (7/10) total participants. Aside from the patient, the other participants were either laboratory members or family members. Conversations were recorded with a directed microphone (Blue Yeti) pointed at the patient and a lavalier microphone (WAYVOSE) on the lapel of the speech partner. All conversation audio streams were recorded directly into the Blackrock recording system, resulting in temporal alignment between speech and neural activity.

**Figure 1.**
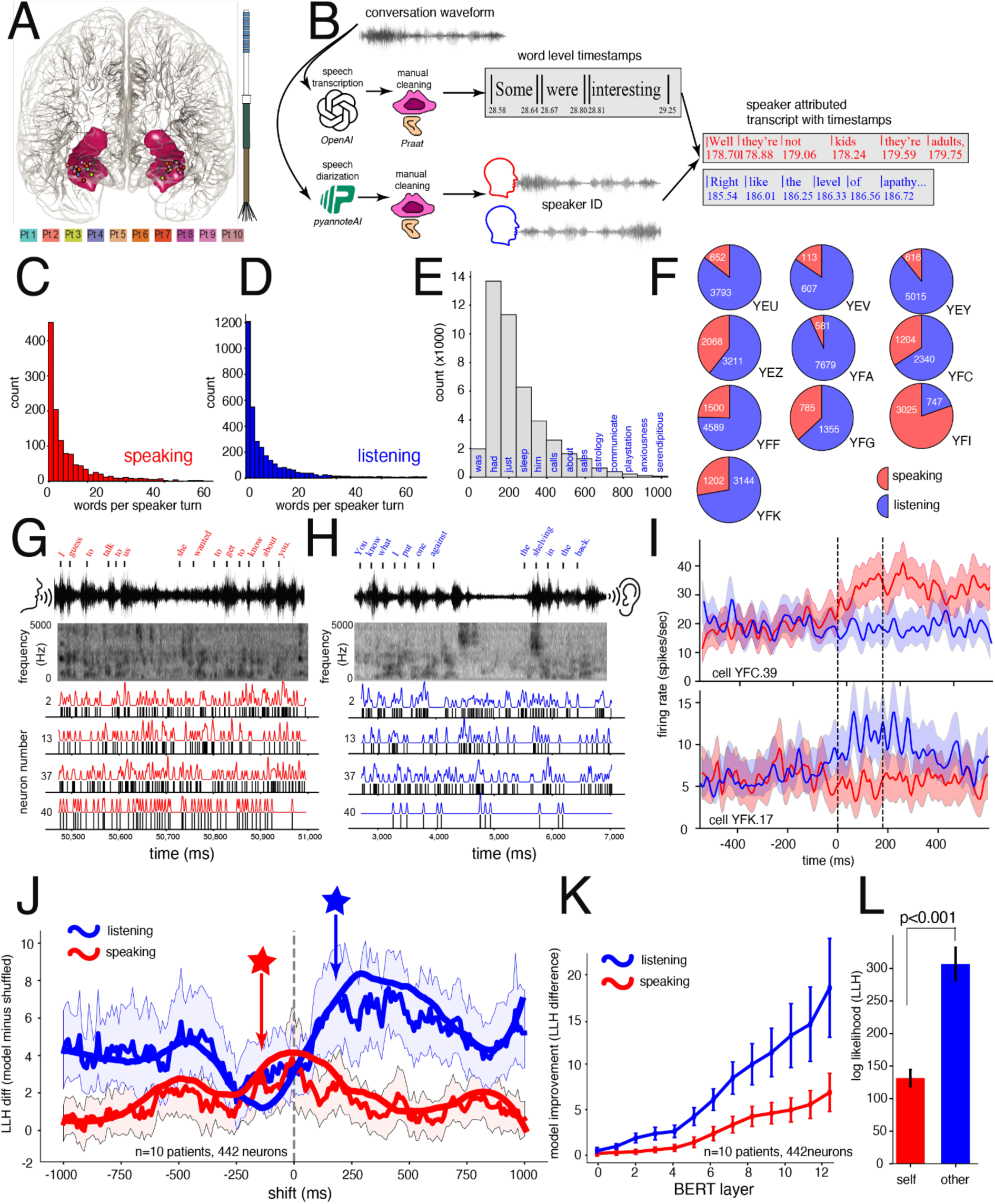
Encoding of semantics of conversational spoken and heard words in the hippocampus. **A.** Recording locations of hippocampal neurons across ten patients. **B.** Schematic of the speech processing pipeline, from raw audio to millisecond-resolved, speaker-labeled transcripts. In this and future panels, red: speaking; blue: listening. **C.** Conversational statistics: distribution of words spoken per turn. **D.** Words heard by the patient per turn from other speakers. **E.** Average word lengths across patients. **F.** Proportion of words produced by each speaker. **G.** Example hippocampal neuron responses from patient YFF showing continuous spiking activity aligned to spoken words. **H.** Same but with heard words and corresponding spectrograms. **I.** Peristimulus time histograms (PSTHs) illustrating turn-aligned responses in two neurons from patients YFC (top) and YFK (bottom). **J.** Temporal dynamics of significant difference in log likelihood between the real and shuffled model (**Methods)** across word-aligned shifts: semantic encoding information began to peak ∼150 ms before word onset for spoken words (red star) and ∼200 ms after word onset for heard words (blue star) according to smoothed data points. **K.** Semantic encoding strength as a function of BERT embedding layer for spoken (red) and heard (blue) words. **L.** Overall encoding strength of the full model for spoken versus heard words.

To identify speech content, we used a combination of automated transcription and diarization with manual correction (**Figure 1B**). Specifically, we began with speech audio files and passed them to two distinct software systems, a lexical decoder/encoder (such as WhisperX), which identified the words that were spoken and their precise onset and offset times, and, in parallel, a diarizer (Pyannote), which identified who was speaking. However, we found that the quality of transcription and diarization were not accurate enough for neural analysis. Thus, specially trained lab members used Praat, a linguistic annotation software for speech analysis (Boersma and Weenink, 2001) to carefully correct both transcription and diarization outputs. That process took roughly one hour per minute of conversation. Finally, all files were manually re-checked and further fixed by first author AGC.

Conversations in the EMU were typical of those observed in other contexts. For example, conversations consisted of short speech turns consisting of a small number of words (median: 3 words; mean: 9 words for spoken, median: 3 words; mean: 10 words for heard. **Figure 1C and D**). Speakers spoke at a rate of 207-261 words/minute. Word lengths ranged from less than 100 ms to almost a full second (**Figure 1E**). Each participants’ speech corpus consisted of an average of 1120 words (**Figure 1F**) and included 325 unique words per patient. Partners’ speech turns consisted of an average of 3178 words (**Figure 1F**), including 644 unique words shared across speakers. Across all ten conversations, 3013 words were used by both the participant and at least one partner; 178 of these were unique. The most common shared words in participant/partners’ conversations were “I”, “like”, “you”, “the”, and “yeah”.

We collected responses of units in hippocampus (HPC, n=442 units, **Figure 1G and H)** from 10 patients (mean: 44.2 neurons; range:19-77 neurons per patient). Neurons showed a clear phasic response to the onset of individual speech turns (two clear examples shown in **Figure 1I**). However, the majority of word-evoked responses were more complex. To understand these responses, we took a word-based encoding model approach (Franch et al., 2025). Specifically, we used a ridge-regularized Poisson regression to predict word-evoked spike trains from static word embeddings (Word2Vec), including both word duration and semantic/duration interaction terms as predictors (**Methods)**. Because our primary interest was in the timing of word-evoked responses, we used a static embedder to avoid contextual bleeding across words and to isolate word-by-word encoding dynamics. Encoding performance was quantified as the difference in log-likelihood between the full model and a control model in which the semantic embedding predictors were shuffled. This analysis revealed distinct temporal dynamics for speaking and listening (**Figure 1J**). Unsurprisingly, responses for spoken words peaked around the time of utterance (red line) while responses for heard word peaked a few hundred ms later (blue line). Guided by these dynamics, we selected a fixed 500 ms window beginning 150 ms before word onset for spoken words and a 500 ms window beginning 200 ms after word onset for heard words. We chose the speaking-related window with the goals of capturing both preparatory and execution-related activity, and because it had a high decoding response. Subsequent analyses confirmed that all results presented are robust to the selection of analysis window. Indeed, other windows tested with a start of +/- 250 ms of the ones we chose produced qualitatively identical results (data not shown).

Once we selected our time windows, we tested the performance on each layer of a contextual embedding model, BERT (Devlin et al., 2019). We chose BERT because it generates context-dependent word embeddings and can incorporate speaker-aware tokens directly into the tokenization process (Gu et al., 2020). We found stronger encoding with higher layers in BERT, which have been shown to capture increasingly abstract and context dependent semantic representation (Jawahar et al., 2019, **Figure 1K**). Notably, encoding was significantly higher (p<.001, Figure 1L) for heard than spoken words.

### On the relationship between neural encoding of heard and spoken language

Responses of three example neurons to three example words that were spoken and heard multiple times are shown in **Figures 2A, B, and C**. While responses to these words were different for speaking and listening, they reflect just a subset of the diversity of neural responses observed. The average word-evoked firing rate of all neurons for all patients together was nearly identical (t=0.268, p=0.7885). Given robust semantic encoding alongside largely nonspecific single-neuron responses across speaking and listening, we next examined the population-level computations that enable semantic generalization while preserving modality-specific structure, consistent with partially overlapping neural codes.

**Figure 2.**
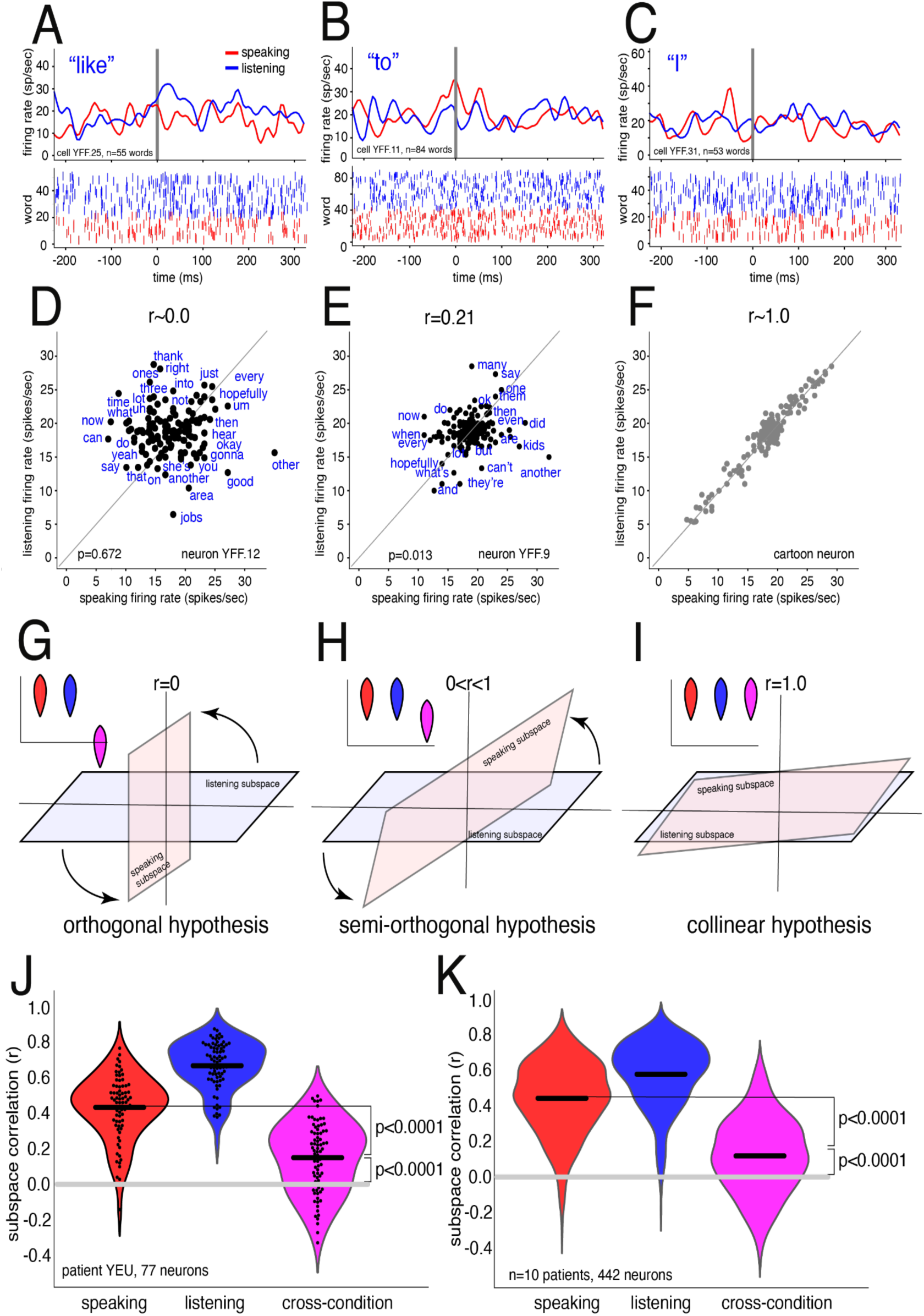
Semi-orthogonality between spoken and heard word representations in the hippocampus. A–C. Example PSTHs and rasters from three hippocampal neurons (patient YFF) during both speaking and listening to the words ***“like”****, **“to”**,* and ***“I”***. These illustrate diverse response patterns across conditions. **D-F** Examples of neuron-level firing rate correlations between spoken and heard words with conceptual schematics: orthogonal/uncorrelated **D.**,semi-orthogonal/partially correlated **E.**, and collinear/perfectly correlated **F. G-I** Competing hypotheses for how hippocampal populations encode word meaning across speaking and listening: orthogonal subspaces or populations with no overlap **G.**, semi-orthogonal subspaces with mixed selectivity across neurons **H.**, and collinear subspaces that are identical across conditions **I. J.** Example within-patient correlations between semantic encoding weights for spoken and heard words: values are significantly above chance but remain below the within-condition half-split noise ceilings. **K.** Group-level results across ten patients show consistent above-chance but sub-ceiling correlations, supporting a **semi orthogonal** mechanism: spoken and heard words are encoded in overlapping but not identical neural subspaces.

We considered three hypotheses for how the same word is represented in neural subspaces for speaking and listening. First, responses to the same word spoken and heard may have no systematic relationship (example neuron, **Figure 2D**). If that pattern were consistent across the population of neurons, then we would expect semantic tuning functions (that is, regression weight vectors) to be uncorrelated. Second, responses may exhibit partial overlap (example neuron, **Figure 2E**), such that tuning functions are positively but imperfectly correlated. Finally, representations may show substantial overlap (cartoon data, **Figure 2F**), in which case tuning correlations would approach their theoretical maximum (in the absence of noise, it would approach 1, although due to noise, it would be lower). These regimes differ in how strongly semantic structure is shared across speakers. At the population level, partial sharing of semantic tuning is consistent with graded orthogonalization, a process that may enable the hippocampus to preserve semantic knowledge while reducing interference between speaking and listening states. Cartoons illustrating the effects of these single neuron solutions on subspace representations are shown in **Figures 2G, H, and I**. These three patterns of encoding correspond to cross-speaker differentiation, differentiation/generalization, and generalization, respectively.

To determine which of these situations our data reflect, we computed the correlation between tuning functions of individual neurons. Here, tuning functions refer to the beta coefficients estimated for each neuron by our ridge regression models fit to spoken versus heard words (**Methods**). Our data are consistent with the intermediate, semi-orthogonal solution (**Figures 2J and K**). Specifically, the degree of orthogonalization between speaking and listening subspaces (purple violins) is greater than zero in all ten of our patients individually (p<0.05 in all cases; one example patient shown in **Figure 2J**), as well as in the group as a whole (p<0.0001, **Figure 2K**). The fact that the measured correlation is greater than zero is, itself, notable since in high dimensional spaces two random vectors are overwhelmingly likely to be orthogonal due to cancellation across dimensions. Instead, this finding indicates that semantic representations are at least partially shared between speaking and listening, offering a solution to the generalization problem.

The degree of overlap observed is less than fully collinear. Of course, an observed correlation less than 1.0 is mathematically guaranteed, but not scientifically interpretable, because any noise from any source will reduce measured correlations below 1.0. The critical question is whether the correlation is less than expected by chance *given the noise properties* of the dataset. We can estimate this using half-split within-condition noise estimates of collinearity. Using these half-split estimates is a powerful control, because it controls against any noise-related process that could cause a spurious positive finding. For example, it may be that fits to the data are not perfect, or that embeddings are suboptimal, or that we chose time windows that are perfectly tailored to our fitting. By comparing real data to half-split within condition data, we control for all of these confounders simultaneously.

We obtained those estimates with both listening (blue violins) and speaking (red violins) data (note that using half-splits reduces our dataset size, thereby reducing correlations, making our analysis even more conservative). We find, both in the population and consistently across participants, lower half-split correlations with speaking than with listening. This fact is consistent with our result, shown above, that overall fits are better for listening than speaking data (**Figure 1L**). For this reason, we take the most conservative approach and focus on the comparison between speaking and speaking/listening cross-condition. We find, for our example patient as well as in the population, that cross-condition correlations are lower than the half-split speaking-only condition (above the null: p=0.0002, below the ceiling p=0.001). We find this same effect in 9/10 of our patients (above the null: p<0.0001, below the ceiling p<0.0001) with the remaining patient showing the same trend (patient YEY, p=0.1124). Together, these results indicate that semantic vectors are poised between orthogonality and collinearity, consistent with the semi-orthogonalization hypothesis for semantic representations in the hippocampus. Note that this analysis does not tell us what drives the differences - how speakers identify speaker identity - but it does tell us that the effects of that identification affect the semantic representation (see **Discussion**).

The observed tuning correlations were not driven by a small subset of neurons. Although 26.47% of individual neurons exhibited significant cross-modality tuning correlations (p < 0.05, assessed against a permutation null obtained by permuting trial correspondence in the listening condition), removing these neurons did not abolish the effect. At the population level, we applied the same permutation procedure and evaluated the mean cross-condition correlation across neurons relative to the corresponding null distribution. The observed mean remained above the permuted null in eight of ten patients, indicating that cross-speaker alignment reflects a distributed population property rather than the contribution of a small group of units.

### Categorical specificity of semantic subspace orthogonality

If two people seated at a table say “pass the salt,” they are likely referring to the same object. In contrast, if one person mentions their eye, they are almost certainly referring to something different from the other’s eye. Different semantic categories, in other words, may have different degrees of cross-speaker generalization and differentiation. We next asked how subspace orthogonalization interacts with semantic categories. We computed categories using a standard approach (Franch et al., 2025; Huth et al., 2016; Jamali et al., 2024; Katlowitz et al., 2025, see **Methods** for details). First, we computed a semantic embedding on all words spoken in all conversations using a non-contextual embedder (Word2Vec, Mikolov et al., 2013) allowing us to capture stable word meanings independent of conversational context. Word embeddings (300-dimensional) were then reduced to two dimensions with UMAP (McInnes et al., 2018), and the resulting space was clustered using HDBSCAN to identify semantic categories in an unsupervised manner. The clustering algorithm yielded 11 semantic categories (**Figure 3A, Methods**). Not surprisingly, function words were the most common (58.6% overall), followed by personal pronouns (13.1% overall). The least used categories were proper nouns, social, and health/medicine (1.3%, 1.6%, and 1.9% of words, **Figure 3B**).

**Figure 3.**
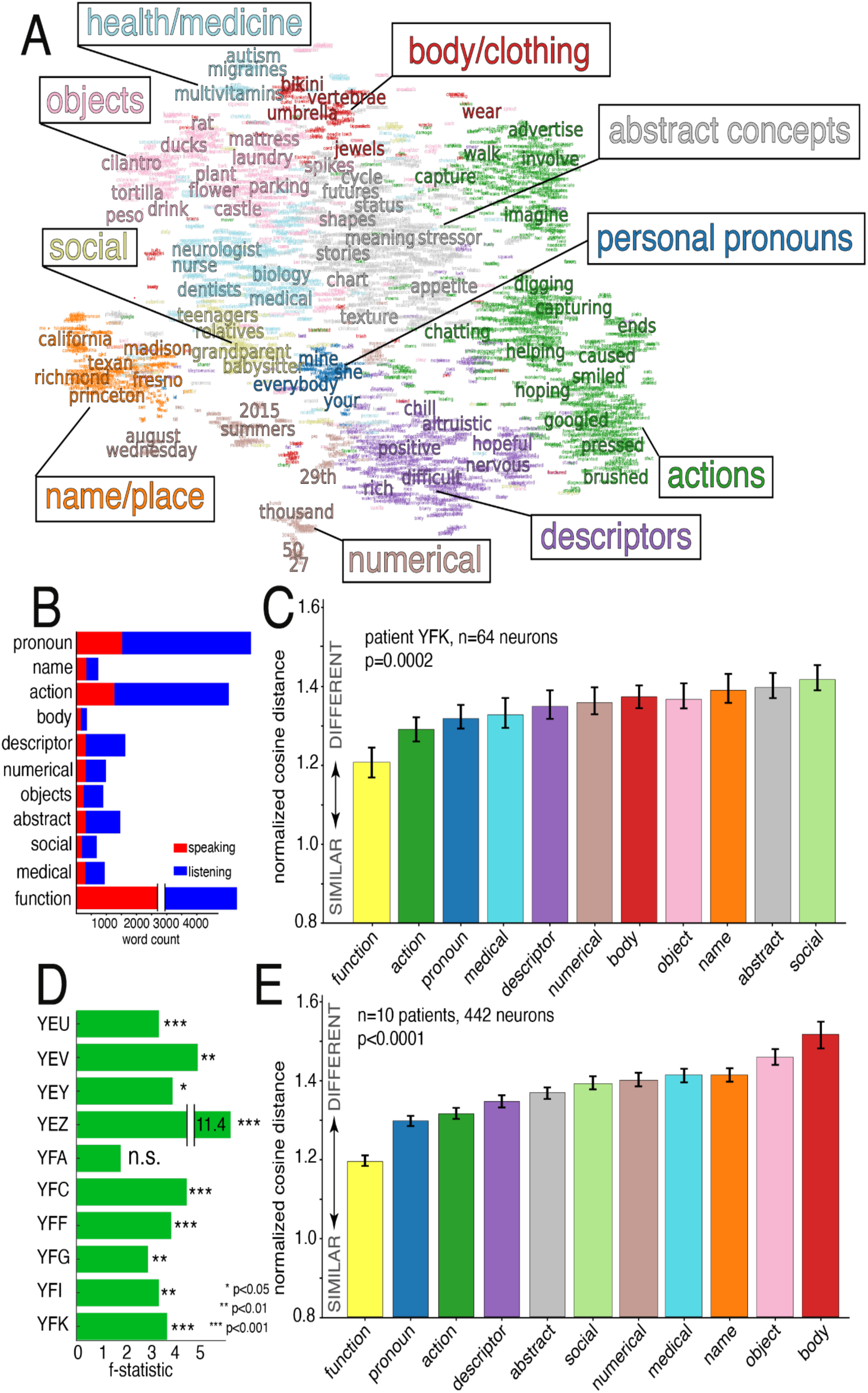
Differential encoding of semantic categories during speaking and listening. **A.** Clustering map of all words produced across the ten patient conversations, revealing broad semantic categories (e.g., body parts, social relationships, function words). **B.** Distribution of words across categories, separated by condition: words produced by the patient (red) versus words spoken by others and therefore heard (blue). Function words were the most common in both conditions. **C.** Example from one patient showing cosine distances between encoding weight vectors for each semantic category, separately for speaking and listening, demonstrating systematic differences across categories. **D.** Patient-level significance summaries: one-way ANOVA tests revealed reliable differences between spoken and heard encoding weights across semantic categories. **E.** Group-averaged cosine distances across patients, highlighting that categories differed in their degree of self–other separation: function words (e.g., *“the,” “and”*) were encoded most similarly across speaking and listening, whereas self-referential categories such as body-related words showed the greatest divergence.

To assess how neural representations of meaning generalize across categories and between speaking and listening, we fit our Poisson ridge regression model (see above and **Methods**) separately within each semantic category, and separately for spoken and heard words. We then measured cosine distance between the resulting tuning vectors. Data from one patient are shown in **Figure 3C**. For this patient, the degree of orthogonality between tuning vectors for self and other varied systematically by semantic category (ANOVA: f-statistic = 3.34; p=0.0002). Function words (example: “*to*”) had the most collinear representations between speaking and listening, followed by action words (“*helping*”) and then, surprisingly, personal pronouns (“*mine*”). Conversely, abstract concepts (“*future*”), social words (“*grandparents*”), and name/places (“*California”)* were represented least similarly between speaking and listening conditions.

This pattern was common across the population. Indeed, of the 10 patients we tested, all but one showed a significant difference across categories (ANOVA, f-statistic =15.64; p<0.0001, **Figure 3D**). Data for the group are shown in **Figure 3E**. Overall, function words were the most similar across participants, followed by personal pronouns and actions. Conversely, body part and clothing terms (“*sweater*”) were the most differentiated, followed by objects and names/places. Together, these results indicate that cross-speaker generalization depends systematically on semantic content. More abstract semantic categories show greater shared representation, whereas more personalized categories exhibit stronger speaker specific differentiation. This pattern indicates that the need to distinguish individualized meanings places constraints on how completely semantic representations can overlap across speakers.

### Hippocampal differentiation of semantic tuning based on conversational partners

Just as we must distinguish our own words from those of our partners, if we have multiple conversational partners, we must keep them separate from each other. If subspace orthogonalization reflects differentiation, rather than low-level differences between articulation and hearing, we would expect similar transformations of semantic representational structure across different non-self conversation partners. Seven of the ten conversations in our dataset had at least three participants. **Figure 4A** shows the number of words spoken by each participant. By convention, we defined the participant as “*speaker 1*” (average of 1120 words), the partner with the most words as “*speaker 2*” (average of 1775 words), and the participant with the next most words as “*speaker 3*” (average of 802 words). We did not analyze data from additional speakers beyond three; in all cases the number of words was too small to meaningfully analyze.

**Figure 4.**
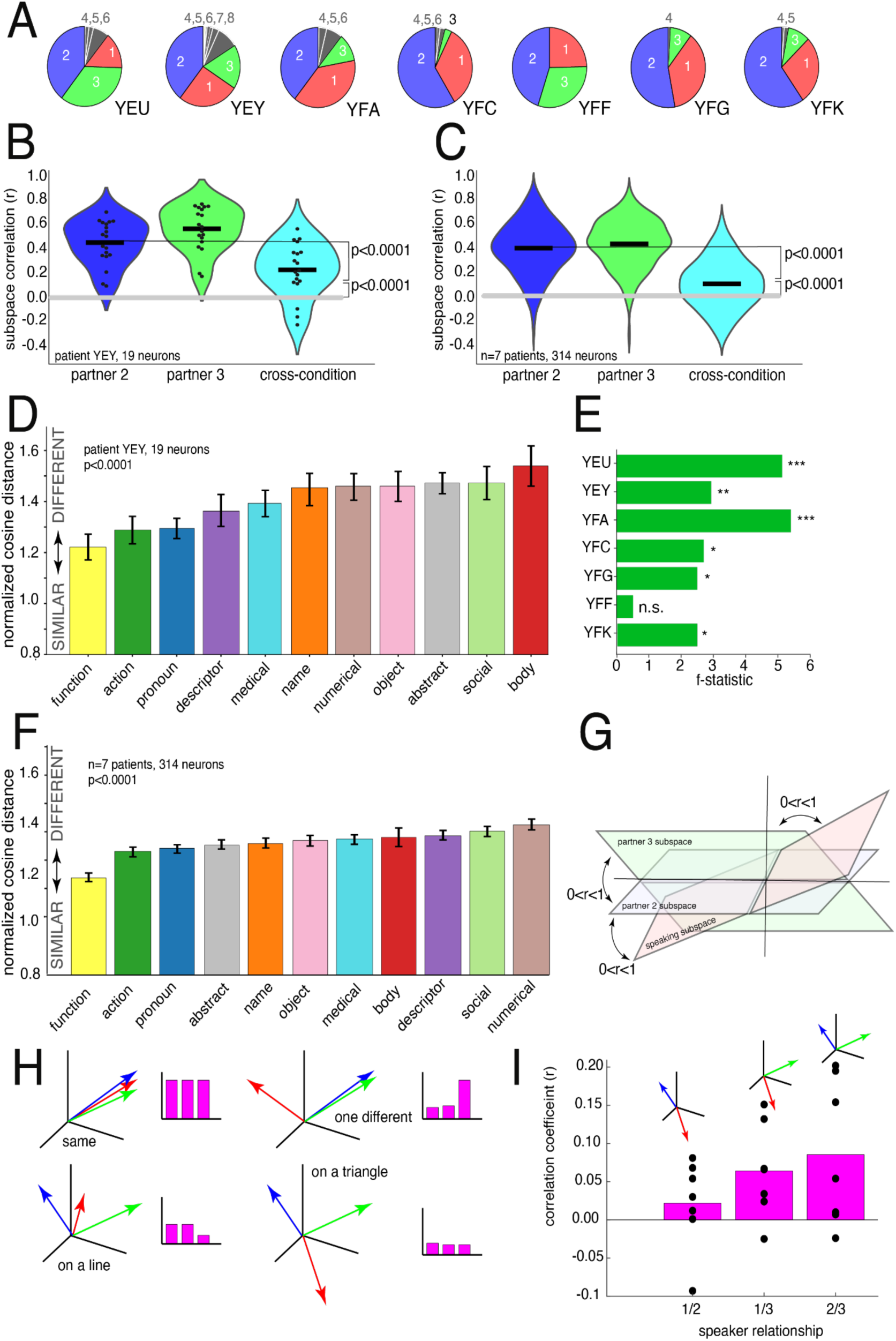
Semantic encoding principles extend beyond self vs. other into differentiation of multiple speakers. **A.** Pie chart of the number of words spoken by unique speakers in each patient’s conversation. Seven patients are highlighted who had more than one other speaker present. **B.** Example patient correlations between semantic encoding weights for words heard from two different speakers. Correlations were significantly above chance but remained below the within-condition half-split noise ceilings. **C.** Group-level results across these seven patients show that the mirror manifold mechanism generalizes to distinguishing between multiple external speakers. **D.** Example from one patient showing cosine distances between encoding weight vectors for each semantic category when listening to two unique speakers. **E.** Patient-level significance summaries: one-way ANOVA tests revealed reliable differences in encoding weights between words spoken by different external speakers. **F.** Group-averaged cosine distances across patients, highlighting that semantic categories varied in their degree of separation: function words were encoded most similarly across speakers, whereas numerical or time-related words were encoded most differently. **G.** Conceptual schematic extending the mirror manifold hypothesis to multiple speakers. **H.** Competing theoretical models for how semantic encoding might scale across speakers: (i) all speakers share the same manifold, (ii) only self versus other are distinct, (iii) external speakers are differentiated but anchored to a common manifold, or (iv) each speaker is encoded on a unique manifold. **I.** Correlation data from all seven patients support the final model: hippocampal neurons encode meaning in a speaker-dependent manner, such that semantic representations are yoked to speaker identity, and each speaker occupies a distinct manifold.

The key finding is that we observe subspace rotation for semantic encodings of partners 2 and 3 (**Figure 4B and C**). Specifically, the subspace correlation between semantics for partner 2 and 3 is greater than zero, indicating that there is some overlap in representations, consistent with the need for cross-speaker semantic generalization (for example patient YEY, p=0.0002; for the population: p<0.0001). However, this overlap was consistently lower than within-condition noise ceilings (patient YEY: p = 0.001; population: p < 0.0001), suggesting that common meanings are expressed in speaker-dependent neural coordinates. As above, using encoding models allows us to isolate effects of semantics from potential low-level confounders such as the speaker’s pitch, tone, distance to the listener, etc (see **Discussion** for more).

We next used the same semantic clustering (**Figure 3A**) to assess semantic category-specific orthogonalization across speech partners. As with self-other contrast, we found a consistent pattern. In both our example participant (**Figure 4D**) and in six of seven participants individually, (**Figure 4E**), we found an effect of semantic category on subspace angle (ANOVA f-stat: 2.93, p = 0.0018). These patterns were also observed when combining across all patients (**Figure 4F**, f-stat: 6.46, p<0.0001). As with the self-other distinction, function words, action words, and pronouns had the most collinearized representations across contexts. In contrast to the self-other distinction, names/titles, objects, and body/clothes were the most distinct for the self-other distinction but were middle positions for the partner 2 vs partner 3 distinction. And, whereas descriptors (“*difficult*”), social (“*grandparents*”), and numerical (*“thousand*”) terms were in the middle for self-other, they were maximally differentiated for partner 2 and partner 3.

Together, these findings suggest that identity is encoded through overlapping semantic representations anchored to a shared backbone and differentiated along speaker-specific axes. The presence of three subspaces (self and two partners) raises the question of how all three are related. All three subspaces may have some degree of orthogonality relative to each other (**Figure 4G**). More generally, we can imagine four possible configurations of three subspaces (**Figure 4H**). They can all be (i) identical, (ii) two matched and one separated, (iii) all three separated, but organized in a line, or (iv) all three separated, forming the points of a triangle. We find the fourth possible configuration, all three separated and forming the points of a triangle (and therefore, consistent with **Figure 4G**). That is, the separating hyperplane that distinguishes self from other is not unique but generalizes to also separate between different others. Thus, we observe not only reliable separation of self from each other speaker, but also measurable separation among the other speakers themselves. Separation is strongest between self and other speakers, although the degree varies depending on which pair is compared. In contrast, separation between different “other” speakers is present but weaker and demonstrated by the higher correlations (**Figure 4I)**. Specifically, correlations for self/other (1/2 and 1/3) are significantly lower than correlations for other/other (2/3, p<0.01, t-test across subjects). This same pattern is observed in 6/7 participants individually (p<0.05 in those cases). In other words, the hippocampus differentiates conversation partners through graded separation of overlapping semantic representations, with stronger differentiation for self versus others than between non-self partners. These findings suggest that hippocampal semantic encoding is organized around a shared semantic structure that is modulated by speaker identity, with the strongest modulation for self-related processing.

### How hippocampal population geometry supports structured and abstract semantic representations

To characterize the hippocampal population geometry associated with cross-modality generalization of meaning, we analyzed the geometry of hippocampal population responses for shared words between listening and speaking. Similar semantic tuning does not necessarily imply preservation of relational structure (Kriegeskorte and Wei, 2021), as neural populations may weight semantic embedding dimensions similarly while organizing word representations differently. Because semantic meaning is inherently relational, preservation of inter-word structure is critical for generalization (Harris, 1954; Firth, 1957; Mikolov et al., 2013). We took the neural population response to words that were both spoken and heard and calculated the pairwise distances to construct a representational dissimilarity matrix (RDM, **Figure 5A**, **Methods**). Representational similarity analysis (Diedrichsen & Kriegeskorte, 2017; Kriegeskorte & Kievit, 2013; Nili et al., 2014) revealed that the geometry of semantics was significantly correlated across speaking and listening within individual patients (permutation test, p<0.0002), **Figure 5B**). For example, this means that if “*somebody*” and “*hurting*” are close in neural space for speaking, then they are close in neural space for listening as well (example see **Figure 5C**). This effect was observed in eight out of ten patients (permutation test, p<0.0002), with weaker effects in two patients likely due to fewer shared words (**Figure 5D**). These findings indicate that the hippocampal neurons construct shared geometry to represent semantics across speaking and listening.

**Figure 5.**
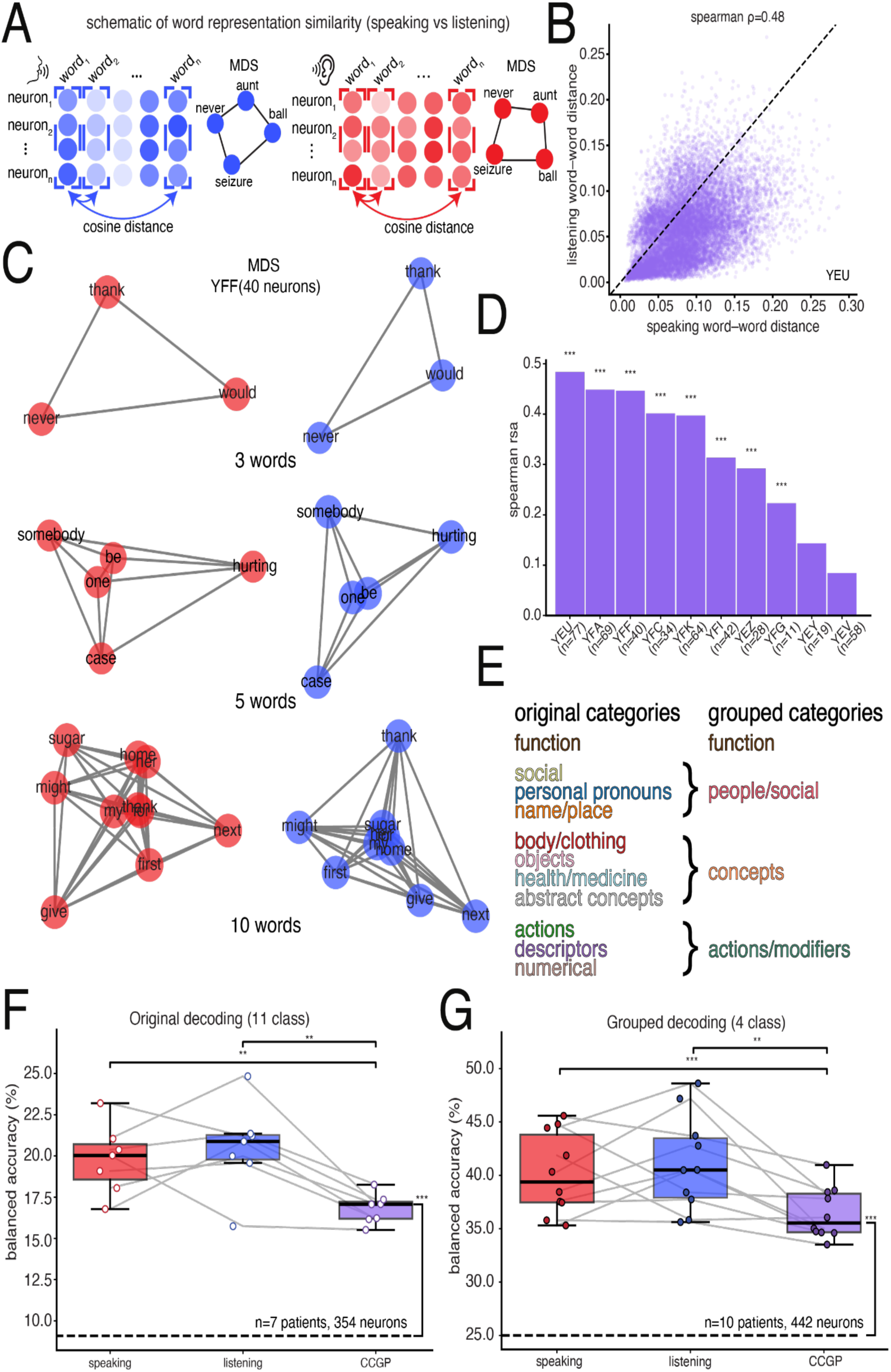
Neural population activity reveals structured and abstract semantic representations across speaking and listening. **A.** Schematic of word evoked neural population responses during speaking and listening to words in common were used to construct representational dissimilarity matrices (RDMs), visualized using multidimensional scaling (MDS) to illustrate the organization of semantic relationships in population space. **B.** Representational similarity analysis for an example patient. Pairwise word–word distances derived from neural population activity during speaking and listening were positively correlated (Spearman ρ). **C.** Multidimensional scaling of speaking and listening RDMs for an example patient across increasing vocabulary sizes. The organization of semantic relationships is preserved locally, while the overall configuration of the representational space differs between conversational roles. **D.** Group-level representational similarity analysis across ten subjects. In eight of ten subjects, correlations between speaking and listening RDMs were significantly greater than those obtained under a null model. **E.** Semantic category grouping used for decoding analyses. Lexical labels were organized into broader conceptual classes to ensure sufficient sampling across subjects and conditions. **F.** Neural population activity supported reliable decoding across eleven lexical categories within both speaking and listening, with cross-condition generalization remaining above chance, consistent with abstraction of semantic information across conversational roles. Each point represents balanced accuracy for an individual subject (n = 7). **G.** Decoding across four broader semantic categories showed reliable within-condition performance and above chance cross condition generalization across all subjects.

The shared semantic geometry across modalities provides a neural substrate for category-level generalization (Nieh et al., 2021, Bernardi et al., 2020), for example, if “*somebody*” and “*hurting*” maintain their proximity in neural space whether heard or spoken, the hippocampus must be encoding abstract meaning rather than modality-specific sensory features. To test this possibility, we assessed cross-conditional generalization performance (CCGP) by training a linear decoder on speaking data and testing on listening data (Bernardi et al., 2020; Courellis et al., 2024). Successful CCGP indicates that the neural population space represents the semantic geometry along modality-invariant axes (Bernardi et al., 2020). We first evaluated decoding across the original eleven semantic categories, and then pooled semantically related categories into four broader groups to maximize statistical power across subjects (**Figure 5E**). This increased the number of samples per class and enabled more reliable estimation of category-level representations. Using logistic regression as our linear decoder, we identified time windows that maximized decoding performance individually for every patient separately for speaking and listening. We observed above-chance decoding (balanced accuracy > chance level, permutation test p < 0.0002) for both modalities as well as robust CCGP (**Figure 5F**). Importantly, significant abstraction persisted when categories were grouped into broader semantic classes, yielding above-chance cross-modal generalization (balanced accuracy > chance level, permutation test p < 0.0002) (**Figure 5G**). Together, these findings demonstrate that hippocampal populations organize semantic knowledge within a shared neural geometry whose abstract structure enables generalization of meaning across speakers.

## DISCUSSION

People engaging in conversation create a temporary shared world that exists in their minds. Consequently, as a conversation participant goes back and forth between listening and speaking, they need to keep track of the fact that words have consistent referents regardless of whether they are spoken or heard. In other words, the brain needs a store of abstract, cross-personal meaning. Here we show that hippocampal responses are consistent with such a pattern. Specifically, we examined word-aligned responses of populations of hippocampal neurons as participants engaged in natural conversations with open topics. Using encoding models, we estimated semantic tuning of both spoken and heard words. We find that there is a shared geometric structure for the two modes, despite their large differences. These differences include motor and premotor demands engaged during both speech planning and perception, as well as differences in interpretation and attention among others. Notably, this shared geometric structure did not require specialized neurons with matched tuning for speaking and hearing (i.e., mirror neurons), but instead was an emergent result of population geometric properties.

Of course, the demands on a cross-personal semantic hub are more than simply abstraction of meaning. This works on two levels. First, in general, we need to somehow bind meaning with speaker identity. Conversational participants need to know who said what. Moreover, specific words can have different contextualized meanings depending on their speaker. Someone talking about their sister refers to a totally different person depending on who is speaking. On the other hand, two people talking about the Astros are probably talking about the same baseball team. So the neural code for meaning needs to be able to be bound - both globally and locally - to the identity of the speaker. Vectorial codes allow for ready binding via representational subspaces (Johnston et al., 2024, Rigotti et al., 2013, Wakhloo et al., 2026, Ebitz and Hayden, 2021). Consistent with this idea, we find graded orthogonalization in subspace geometry for spoken and heard meaning - and for meaning from different heard conversation partners. Importantly, our use of encoding models derived from speech transcripts means that we can specifically link these subspace changes to semantic features of the words, rather than potential confounders like pitch, prosody, speaker identity, speaker relationship, and so on. Our work, therefore, contributes to a growing body of work suggesting that trade-offs between generalization and differentiation can be supported by structured relationships between neural population subspaces, including graded alignment or partial overlap (Bernardi et al., 2020; Elsayed et al., 2016; Kaufman et al., 2014; Libby and Buschman, 2021; Tang et al., 2020; Xie et al., 2022; Johnston et al., 2024; Tian et al., 2024; Chericoni et al., 2025). These ideas are applicable to structured representations of the hippocampus (Bernardi et al., 2020; Courellis et al., 2024); our work therefore extends these findings to the domain of language.

Our study complements recent work on conversational language. Prior studies show that contextual embeddings from large language models predict neural activity during both production and comprehension with temporally structured dynamics that mirror conversational exchange (Zada et al., 2024; Goldstein et al., 2025). Other work using intracranial recordings demonstrates that natural conversation recruits distributed cortical and limbic networks, with only partial overlap between production and comprehension related activity (Cai et al., 2025). Evidence for bridge and mirror neurons in the precentral cortex further suggests shared responses to speech elements across listening and speaking, although such neurons do not account for flexible speaker-specific representations (Xu et al., 2024). Together, these studies highlight the existence of shared neural substrates for conversational language while leaving unresolved how semantic representations generalize across speakers. Our results extend prior work by demonstrating that cross-speaker representational overlap can be identified in natural, free-flowing speech. First, we show that such overlap emerges under unconstrained conversational conditions. Second, we find that this organization is not limited to phonemic features but extends to semantic representations. Third, we show that semantic structure is represented outside the classical language network. Finally, we demonstrate that shared semantic structure and identity-dependent differentiation coexist within the same neural populations, consistent with a unifying population-level framework in which communication is supported by structured representational geometry rather than specialized neuron types or modality-specific channels.

Our results may be surprising as we find clear semantic encoding localized to the hippocampus, which is not a classical language structure (Fedorenko et al., 2024). However, our results are aligned with a good deal of research highlighting the role of the hippocampus in representing word meanings (Binder et al., 2009; Brown-Schmidt et al., 2021; Duff and Brown-Schmidt 2012; Franch et al., 2025; Piai et al., 2016; van de Ven et al., 2020) and grammatical processing (Dijksterhuis et al., 2024). Complementary studies of lesions to the hippocampus show clear semantic deficits (Manns et al., 2003; Moscovitch et al., 2005; Keane et al., 2019; Duff et al., 2020). This possibility is consistent with the robust evidence that the hippocampus plays an important role in memory-related processes, which may be foundational for semantics. Moreover, the hippocampus is associated with simultaneous generalization and differentiation (Bernardi et al., 2020), as well as both pattern completion and separation processes, which are conceptually related (Leutgeb et al., 2005; Bakker et al., 2008). Finally, there is evidence that supports a potential role for the hippocampus in social cognition, including identity assignment (Danjo et al., 2018; Tavares et al., 2015; Montagrin et al., 2018; Rey et al., 2020).

The semantic clustering we observe indicates that subspace rotation is not a one-size-fits-all process (**Figure 3**). While our results do not provide insight into why certain categories have more or less alignment, we can speculate. For example, function words (*the*, for, *and*) and pronouns are the most consistent across both self-other (**Figure 3E**) and partner2-partner3 (**Figure 4F**). These words have both very general meanings that do not vary much by person and, moreover, tend to have very weak semantics on their own (Katlowitz et al., 2025). Verbs are the next most consistent (*helping*, *brushed*, *smiled*). Conversely, body parts (*face*, *eyes*, *arm*) tend to be highly specific to individuals; the speakers’ hand is almost certain to be different - and have different conversational relevance - than the partner’s hand. That may also explain why objects (*drink*, *laundry*, *mattress*) are highly differentiated - speakers may tend to discuss objects relative to themselves.

Some recent functional-anatomical accounts emphasize that production and comprehension are carried out in partially separable hubs or dual processing streams (e.g., Hickok & Poeppel, 2007; Ben-Shalom & Poeppel, 2008; Castellucci et al., 2022). Such dichotomous frameworks leave unresolved the need for cross-speaker generalization. This problem motivates the development of perspectives that suggest that speaking and listening may draw on shared representational systems (Pickering & Garrod, 2013; Tremblay & Dick, 2016; Silbert et al., 2014). Our results build on these ideas and further suggest that this overlap is structured in a manner that preserves shared semantic organization while allowing identity-dependent differentiation. More broadly, our findings suggest that the representational principles supporting cross-speaker generalization of meaning may extend to other domains requiring self–other differentiation, such as mind-reading and mental state inference (Mukamel et al., 2010; Koster-Hale and Saxe, 2013; Jamali et al., 2021). As in conversational language, these processes require simultaneously recognizing shared structure between self and other while maintaining their separation (Uddin, 2011; Jamali et al., 2021; Heatherton et al., 2016). For example, empathy depends on representing another person’s experience while preserving its distinction from one’s own (Decety and Jackson, 2004; Singer et al., 2004; Lamm et al., 2007). In this context, our results suggest that semantic information is encoded as an abstract variable within hippocampal population activity, supporting generalization across individuals while preserving identity-specific distinctions. That we observe such an organization in the context of natural conversation suggests it may provide a foundation for broader cognitive functions, including emotion, theory of mind, and social decision-making.

## METHODS

### Human intracranial neurophysiology

All experiments were performed in the Epilepsy Monitoring Unit (EMU) at Baylor St. Luke’s Hospital using standardized approaches (Xiao et al., 2024). Experimental data were recorded from ten adult patients (seven male, three female, age ranges: 22-54) undergoing intracranial monitoring for epilepsy. The hippocampus was not a seizure focus area of any patients included in the study. Single neuron data were recorded from stereotactic (sEEG) probes, specifically Ad-Tech Medical probes in a Behnke-Fried configuration. Each patient had an average of three probes terminating in the left and right hippocampus. Electrode locations were verified by co-registered pre-operative MRI and post-operative CT scans. Each probe includes eight microwires, each with eight contacts, specifically designed for recording single-neuron activity. Single neuron data were recorded using a 512-channel *Blackrock Microsystems Neuroport* system sampled at 30 kHz. To identify single neuron action potentials, the raw traces were spike sorted using the *Wave_clus* sorting algorithm (Chaure et al., 2018) and then manually evaluated and modified to improve sorting. Noise was removed and each signal was classified as multi or single unit using several criteria: consistent spike waveforms, waveform shape (slope, amplitude, trough-to-peak), and exponentially decaying ISI histogram with refractory period (1 ms) violations below 5%. The analyses here used all single and multiunit activity.

### Electrode visualization

Electrodes were localized using the software pipeline intracranial Electrode Visualization (iELVis; Groppe et al., 2017) and plotted across patients on an average brain using Reproducible Analysis & Visualization of iEEG (RAVE; Magnotti et al., 2020). For each patient, DICOM images of the preoperative T1 anatomical MRI and the postoperative Stealth CT scans were acquired and converted to NIfTI format (Li et al., 2016). The CT was aligned to MRI space using FSL (Jenkinson et al., 2002; Jenkinson & Smith, 2001). The resulting coregistered CT was loaded into BioImage Suite (version 3.5β1; Joshi et al., 2011) and the electrode contacts were manually localized. Electrodes coordinates were converted to patient native space using iELVis MATLAB functions (Yang et al., 2012) and plotted on the Freesurfer (version 7.4.1) reconstructed brain surface (Dale et al., 1999). Microelectrode coordinates are taken from the first (deepest) macro contact on the Ad-Tech Behnke Fried depth electrodes. RAVE (Magnotti et al., 2020) was used to transform each patient’s brain and electrode coordinates into MNI152 average space. The average coordinates were plotted together on a glass brain with the hippocampus segmentation and colored by patient.

### Conversations

Patients with epilepsy and healthy language function engaged in unprompted conversations with researchers, family members, friends, and/or members of the clinical staff. Conversation lengths ranged from 14.7-70.1 minutes, with an average length of 20.2 minutes. Topics of conversation varied across patients. Audio was primarily recorded using two room microphones that were directed toward the location of the patient (Blue Yeti) or personal lapel microphones (WAYVOSE). The audio was synchronized to the neural recording system via analog input going directly into the Blackrock Neural Signal Processor at 30 kHz.

### Audio Transcription

After experiments, the audio.wav file was automatically transcribed using WhisperX, a state-of-the-art and commercially available AI model trained to transcribe speech (Bain et al., 2023). Speaker diarization was achieved using an open-source toolkit written in Python named Pyannote (Bredin et al., 2021). The transcribed words, corresponding timestamps, and speaker labels output from these models was converted to a TextGrid and then loaded into Praat, a software for speech analysis. The original.wav file was also loaded into Praat. Trained lab members manually inspected the spectrograms, timestamps, and speaker labels, correcting each word to ensure that onset and offset times were precise and assigned to the correct speaker (this process took about 1 hour per minute of speech). The TextGrid output of corrected words and timestamps from Praat was converted to a.xlsx and loaded in Python for further analysis.

### Contextual and non-contextual embeddings

#### Contextual embeddings

To obtain contextual embeddings for each word, we used the *bert-base-cased* model (BERT; Devlin et al., 2019) accessed via the Hugging Face Transformers library (Wolf et al., 2020). This model consists of one embedding layer and 12 transformer blocks, each producing a 768-dimensional vector for every token. The conversational transcripts were vectorized into a single sequence of words in their natural temporal order and each word was prepended with a speaker-specific token, allowing the model to differentiate between speakers. This modification was inspired by prior speaker-aware adaptations of BERT for dialogue modeling (Gu et al., 2021). To handle BERT’s 512-token input limit, we used a sliding window of 512 tokens with a stride of 256, and averaged embeddings for words that fell in overlapping window segments. This model also allows for extraction of embeddings from each layer (0-12). Embeddings from each of these layers were used as predictors for neural activity (**Figure 1K**), however all other results are from output or final layer.

#### Non-contextual (static) embeddings

We also extracted static word embeddings using the pre-trained fastText Word2Vec model in MATLAB, specifically the wiki-news-300d-1M vectors (Bojanowski et al., 2017; Grave et al., 2018). This model provides fixed 300-dimensional vectors for over one million words and phrases, trained on Wikipedia and news text to capture broad semantic relationships. Unlike contextual embeddings, which vary depending on sentence position or surrounding words, these static embeddings assign the same representation to each word regardless of context. For our dataset, we mapped every word in the transcripts to its corresponding fastText vector. Words without valid embeddings (e.g., uncommon surnames or rare tokens not present in the model’s vocabulary) were excluded from further analyses.

#### Semantic clustering

To identify natural semantic categories in our word data (**Figure 3A–E**), we clustered all unique words into groups based on distributional similarity in embedding space. We first extracted 300-dimensional static embeddings for each unique word across all conversations using the pre-trained fastText wiki-news-300d-1M model. To reduce noise and focus on semantically informative tokens, we excluded function words, except personal pronouns and their contractions, using the stopword list from the Natural Language Toolkit (NLTK; Bird et al., 2009), and treated them as a separate cluster. Dimensionality reduction was then performed with Uniform Manifold Approximation and Projection (UMAP; McInnes et al., 2018) using the parameters *n_neighbors = 15*, *min_dist = 0.2*, *n_components = 10*, and *metric = cosine*. UMAP produces a lower-dimensional representation in which words with similar meanings are located nearby. We next applied Hierarchical Density-Based Spatial Clustering of Applications with Noise (HDBSCAN; Campello et al., 2013; McInnes & Healy, 2017) to these reduced embeddings. HDBSCAN requires two key parameters: *minimum cluster size* (the smallest grouping to be considered a valid cluster) and *minimum samples* (the number of points required to define a dense region), which together determine the stability and granularity of the resulting clusters. In our analysis, we set *minimum cluster size = 40* and *minimum samples = 8*. Words assigned to the noise cluster were reassigned to the nearest cluster by majority vote of their neighbors. For visualization, we projected the raw embeddings into two dimensions using t-distributed Stochastic Neighbor Embedding (t-SNE; van der Maaten & Hinton, 2008) and colored words by their assigned cluster IDs.

#### Turn-aligned PSTHs and rasters

For each neuron, we extracted spike trains aligned to conversational turn onsets defined as the moment when either the patient (self) began speaking or when any other participant in the room initiated speech (listening for self). Each turn onset was treated as time zero, and spikes were segmented into fixed analysis windows spanning a defined pre-and post-onset interval. Within each window, spikes were binned at a fixed bin width (20 ms). Spike counts in each bin were divided by the bin duration (in seconds) to yield firing rates in Hz and averaged across trials to obtain the mean peri-event time course of firing rate (PSTH). Variability across trials was quantified as the standard error of the mean (SEM). For raster visualization, individual spikes from each trial were plotted as vertical ticks at their millisecond timestamps relative to turn onset, forming rasters at the baseline of the axis. To highlight temporal structure, the binary spike trains were also smoothed with a Gaussian kernel (σ ≈ 10 ms) and scaled to approximate firing rate. These smoothed traces were plotted directly above the raster ticks, providing a continuous PSTH trace superimposed on the underlying spike times.

### Semantic Encoding Analyses

#### Regressing spike counts on word embeddings

Word embeddings, such as those derived from Word2Vec or similar models, are typically high-dimensional and can exhibit substantial collinearity among dimensions - that is, many of the embedding features are correlated with one another. We first used Principal Component Analysis (PCA, scikitlearn) on the full word embeddings to obtain uncorrelated features that still capture the dominant structure in the embedding space with reduced dimensionality. For each word, we used the first 100 principal components (PC) from Word2Vec or BERT embeddings because they explained at least 60% of the variance of the original embedding vectors. We modeled the spike count responses of individual neurons using a Poisson Generalized Linear Model (GLM) with a log-link function and ridge (L2) regularization (we confirmed that spike counts for each word follow a Poisson distribution). The model aimed to predict the number of spikes a neuron fired in response to each word (summed spikes across word duration), using the top 100 principal components (PCs) of the word embedding vector, the duration of the word, and the interaction between each PC and the word’s duration as predictors (z-scored prior to model fitting). This resulted in a 201-dimensional feature vector for each word: 100 PC values, 1 duration value, and 100 PC×duration interaction terms.

The Poisson GLM assumes that the spike count *y_i_* for word *i* is drawn from a Poisson distribution with mean λᵢ where:

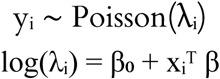

Where,the expected spike count for word (*i)* and neuron (*n*), is modeled by the linear regression coefficients (β), one for each of the 201 predictors, 100 PCs from embeddings, word duration, and duration PC interactions, (X_i_^T^), plus the y-intercept (β₀). For each neuron, we used a 5-iteration nested training loop, where in each iteration, the data was split into training and held-out test sets (80/20 split). Within the training set, 5-fold cross-validation was used to select the optimal regularization strength (alpha) from a log-spaced range based on optimizing the cross-validated log-likelihood. The selected model was then evaluated on the held-out test set using several performance metrics, including log-likelihood, and adjusted R². To determine whether the observed performance exceeded chance, we fit matched null models in which embeddings were randomly shuffled across words 100 times, computing LLH from every shuffled model. To determine if the actual model showed significant improvement from the shuffled model, we performed a permutation test comparing the actual model LLH to the shuffled model LLH distribution. We then took the difference between the actual and shuffled model (median) LLH (LLH diff; ΔLLH). A neuron’s response was successfully predicted by the model if its ΔLLH was both positive and significant and if McFadden’s pseudo-R² value was positive (p < 0.05). Shuffled model R² values were significantly less than those from actual models (p < 0.05, Wilcoxon signed-rank test). A predictor was considered significant if its corresponding coefficient had a p-value less than 0.05 in the fitted Poisson model. For population-level encoding analyses, however, we included all neurons and all model coefficients, irrespective of their individual significance.

#### Hyperparameter Selection

To choose the regularization strength α, we used a two-stage grid search:

1. **Coarse Search** over:^αɛ{10-5, 10-4,…,103}^
2. **Fine Search** around the best coarse value, spanning two orders of magnitude.

For each value of *α*, we performed 5-fold cross-validation. For each fold, the model was trained on the other four folds and evaluated on the held-out fold by computing the Poisson log-likelihood:

#### Window selection for speech comprehension and production

To identify the analysis windows for subsequent encoding models, we systematically quantified model performance across a range of candidate window starts and lengths. Candidate windows were defined from –1 s to +1 s relative to word onset, sampled in 10-ms increments, with window lengths ranging from 80 ms to 500 ms. For each [window start × window length] combination, and separately for spoken and heard words, we fit Poisson regression models using Word2Vec embeddings as predictors. Performance was evaluated using ΔLLH, defined as the log-likelihood difference between the full model and a semantic-shuffled model in which only the semantic embedding PCs were permuted (with duration interactions recomputed). Model performance increased monotonically with longer window lengths, with the strongest improvements at 500 ms. Based on this, and to capture preparatory activity prior to speech and integrative activity during comprehension, we selected window starts that were shifted back from the global maximum. For speaking, the window began 150 ms before word onset; for listening, the window began 200 ms after word onset. For visualization, ΔLLH values were smoothed across window positions using a Savitzky–Golay filter (Savitzky & Golay, 1964). For each neuron and word, firing rate was computed as the total spike count within the selected window, divided by the window duration, and expressed in spikes/s. Unless otherwise noted, all analyses reported in the manuscript used these fixed windows (**Figures 1I and 2A–C** show PSTHs computed using identical windows for speaking and listening).

#### Reliability and cross-condition comparisons of semantic encoding

To evaluate how neural populations encode semantic information (using BERT embeddings) across speaking and listening, we compared the structure of regression weights obtained under each condition. We fit separate Poisson ridge regression models for spoken and heard words (see above), yielding condition-specific vectors of beta coefficients that define each neuron’s semantic tuning curve. However, to dismiss possible explanations of these results coming from an imbalance in the number of words within each condition we made sure to downsample the larger class if the imbalance exceeded 1.5 times the smaller class as implemented in the imbalanced-learn Python package (Lemaitre et al., 2017). These results did not vary significantly from refitting the same models on perfectly balanced data. In our framework, the regression weights learned by the model can be interpreted as the tuning curves. When considered jointly across neurons, these tuning vectors span a neural subspace that captures the population-level encoding of semantics under a given condition (speaking or listening). The alignment of these two subspaces was quantified as the correlation between their coefficient vectors (*r*_cross) (Johnston et al., 2024). To assess whether these correlations exceeded chance levels, we constructed a null distribution by permuting the predictor variables: one condition was modeled on the true data, while the other was modeled on data with semantic predictors randomly shuffled. This procedure was repeated 500 times to estimate the distribution of correlations expected under the null hypothesis of no shared structure. As an upper bound, or noise ceiling, we quantified within-condition reliability by randomly splitting the data in half and re-fitting the model separately to each split. Correlations between the resulting beta vectors provided half-split reliability estimates for spoken (*r*_self) and heard (*r*_other) words. Together, these three metrics allowed us to determine whether cross-condition similarity reflected true shared semantic encoding rather than noise, and to interpret it relative to the maximum reliability attainable in each condition.

#### Semantic cosine distance

We quantified the separation between semantic encoding during speaking and listening using the cosine distance between regression weight vectors. For two beta coefficient vectors w_1_ and w_2_ cosine distance is defined as:

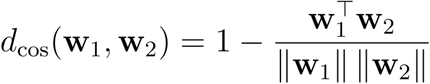

To compute category-specific distances, the regression model described above was fit separately for each semantic category under speaking and listening conditions. This procedure was repeated across all categories, with class counts balanced up to an imbalance ratio of 1.5 to avoid excessive trial loss. To establish a noise ceiling for these distances, we computed half-split reliability estimates within the most populated category (function words). Data from this category were randomly split in half, models were refit, and cosine distances were computed between the two halves. This was repeated with bootstrapping, and all observed category distances were normalized by the mean half-split distance for function words. Finally, differences in normalized cosine distances across semantic categories were evaluated using a one-way ANOVA.

### Geometric similarity analyses

#### Construction of Representational Dissimilarity Matrices (RDM)

For both speaking and listening conditions, neural representational dissimilarity matrices (RDMs) were constructed from population response patterns using cosine distance as the dissimilarity metric (Diedrichsen & Kriegeskorte, 2017; Kriegeskorte & Kievit, 2013; Nili et al., 2014). Neural responses were first organized into neuron-by-word matrices Χ_ℓ_∈ℝ^p×n^ where *p* denotes the number of recorded neurons, and *n* denotes the number of lexical items shared across conditions, and each column **r***_i_*^(*l*)^ represents the population response vector evoked byword i in condition ℓ, corresponding to **r***_i_*^(*l*)^. Repeated occurrences of the same word were averaged prior to analysis to obtain a single population vector per lexical item.

The neural RDM for condition was defined as the matrix of pairwise cosine distances between all word-evoked population vectors:

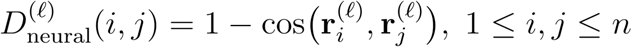

This procedure yields a symmetric n×n dissimilarity matrix with zero diagonal. For downstream analyses, each RDM was vectorized by extracting the upper triangular elements, resulting in a condensed dissimilarity vector of length 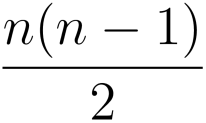 containing all unique pairwise distances.

#### Representation similarity analyses

To quantify the similarity of neural representational geometries across conversational roles, we performed representational similarity analysis (RSA) between speaking and listening neural RDMs. For each condition, we extracted the upper triangular elements of the RDM (excluding the diagonal) to form condensed dissimilarity vectors capturing all unique pairwise relationships between word-evoked neural responses. Cross-condition geometry similarity was quantified as the Spearman rank correlation (Nili et al., 2014) between the vectorized speaking and listening RDMs. A high positive correlation indicates that word pairs that are represented similarly in one condition (i.e., small neural distance) are also represented similarly in the other, reflecting a shared neural geometry across conversational roles.

To assess statistical significance, we generated a null distribution using a permutation test. The speaking RDM was held fixed, while the listening RDM was jointly permuted along its rows and columns using the same random permutation of word identities. This procedure preserves the intrinsic representational geometry within the listening condition while disrupting correspondence between conditions. For each permutation, the upper-triangular dissimilarity vector of the permuted listening RDM was recomputed and correlated with the fixed speaking vector to generate a null distribution of correlation values. The p-value was computed as the proportion of permuted correlations exceeding the observed value. RSA was computed separately for each patient.

#### Multidimensional Scaling (MDS) and network visualization

We employed classical Multidimensional Scaling (MDS) (Mead, 1992) combined with network visualization techniques to visualize the geometric organization of neural representations for matched word pairs across languages. This approach reveals the topological structure of semantic representations in neural space by projecting high-dimensional neural patterns onto interpretable low-dimensional manifolds while preserving pairwise dissimilarities. The resulting 2D coordinates were visualized as network graphs where nodes represent words and weighted edges encode distance between words.

### Decoding Analyses

#### Temporal alignment and feature construction for decoding

Neural firing rates were estimated by convolving spike trains with a Gaussian kernel (σ = 20 ms) to obtain a continuous firing rate signal, which was then discretized into 20 ms bins. To align neural activity to word events while maintaining comparability across trials of varying duration, we constructed time windows anchored to word onset and offset. For each word, neural activity was extracted from a window spanning the word duration, with additional temporal padding applied before and/or after the word to ensure a fixed window length of 500 ms across all trials.

To account for uncertainty in the temporal alignment between neural responses and linguistic events, we systematically varied the onset of the analysis window relative to each word (−500 to 500 ms). This temporal sweep allowed us to identify the time period that maximized decoding performance separately for speaking and listening conditions. For each window configuration, firing rates were averaged within the window to yield a single population response vector per word.

#### Semantic category linear decoder and cross condition generalization performance

Semantic category decoding was performed using a multinomial logistic regression classifier with ℓ2 regularization. Model hyperparameters (regularization strength) were selected via three-fold cross-validation with three repeated splits to ensure stability of model performance estimates.

To mitigate class imbalance across semantic categories, we applied a balancing procedure such that categories contributed approximately equally to the training data. This ensured that decoding performance was not driven by overrepresented categories and enabled a more uniform assessment of category-level encoding.

Decoding performance was quantified using balanced accuracy. For each patient, we first identified the temporal window that maximized within-condition decoding performance separately for speaking and listening conditions. These condition-specific optimal windows were then used to evaluate cross-condition generalization performance.

Cross-condition generalization (CCGP, Bernardi et al., 2020) was assessed by training the decoder on neural responses from one condition (e.g., speaking) and testing on the other (e.g., listening), and vice versa, the average of the two was used as the final CCGP metric. This approach quantifies the extent to which semantic representations learned in one conversational role generalize to the other.

To assess statistical significance of decoding performance, we generated null distributions using permutation testing. Specifically, semantic category labels were randomly shuffled within the training data, and the decoding procedure was repeated using the same cross-validation and evaluation pipeline. The p-value for each decoding analysis was computed as the proportion of permuted decoding accuracies that exceeded the observed balanced accuracy. Paired t-tests were used to assess whether decoding performance differed between within-condition decoding to cross condition generalization performance.

## Acknowledgements

We thank Joshua Adkinson, Justin Fine, Victoria Gates, Suzanne Kemmer, George Kokalas, and Steven Piantadosi for invaluable assistance.

